# Avoiding attack: How dune wasps leverage colour and motion to detect their cryptic spider predators

**DOI:** 10.1101/2021.03.20.436281

**Authors:** Dulce Rodríguez-Morales, Horacio Tapia-McClung, Luis Robledo-Ospina, Dinesh Rao

## Abstract

Ambush predators depend on cryptic body colouration, stillness and a suitable hunting location to optimise the probability of prey capture. Detection of cryptic predators, such as crab spiders, by flower seeking wasps may also be hindered by wind induced movement of the flowers themselves. In a beach dune habitat, *Microbembex nigrifrons* wasps approaching flowerheads of the *Palafoxia lindenii* plant need to evaluate the flowers to avoid spider attack. Wasps may detect spiders through colour and movement cues. We tracked the flight trajectories of dune wasps as they approached occupied and unoccupied flowers under two movement conditions; when the flowers were still or moving. We simulated the appearance of the spider and the flower using psychophysical visual modelling techniques and related it to the decisions made by the wasp to land or avoid the flower. Wasps could discriminate spiders only at a very close range, and this was reflected in the shape of their trajectories. Wasps were more prone to making errors in threat assessment when the flowers are moving. Our results suggest that dune wasp predation risk is augmented by abiotic conditions such as wind and compromises their early detection capabilities.

## INTRODUCTION

Prey strategies to avoid attack by ambush predators are more effective in the early part of the predation sequence (Pembury Smith and Ruxton, 2020; Ruxton et al., 2018). Ambush predators are often cryptic, with their body colouration matching their environment (Anderson and Dodson, 2015); they are very still since movement can break their crypsis (Gonzálvez and Rodríguez-Gironés, 2013); they often have venom to debilitate their prey (Schwantes et al., 2018) and most importantly they hunt at a moment when the prey is vulnerable, i.e., during foraging or mating -- when prey awareness is compromised (Pembury Smith and Ruxton, 2020). Therefore, for a prey to overcome an ambush predator’s strategy, evaluation of a risky site is crucial.

A prey’s ability to detect an ambush predator is constrained by its perceptual capabilities either through chemical or visual mechanisms (Gonzálvez and Rodríguez-Gironés, 2013). Visual detection of a predator depends on the spectral sensitivity of the prey’s eye (the ability of the eye to respond to specific wavelengths of the light spectrum; Cronin et al., (2014)), spatial acuity (the capacity to discriminate shape and pattern details; Caves et al., (2018)) and temporal resolution (time taken to process visual information; Cronin et al., (2014)). Furthermore, abiotic factors such as wind or obstacles can add to the visual clutter in a habitat (Burnett et al., 2020; Hennessy et al., 2020) and consequently hinder predator detection.

The problem of detecting ambush predators is commonly faced by pollinating insects that approach flowers harbouring crab spiders (Araneae: Thomisidae) (Morse, 1986). Crab spiders are famously cryptic -- their body colouration blends into the background of the flowers (Thery and Casas, 2009); some species are capable of changing their colour to match the flower (Oxford and Gillespie, 1998) and others are mottled in various shades (Rodríguez-Morales et al., 2018). This crypsis may serve to avoid detection by potential prey (Gavini et al., 2019; Morse, 1986), perhaps by interfering with search image formation. However, there is still controversy whether the intended receivers of the crypsis are prey or predators. Crypsis was found to be ineffective when considering the entire community of flower visiting potential prey (Brechbuhl et al., 2010) and a recent study argued that crypsis in crab spiders reduce their risk of predation by birds (Rodríguez-Gironés and Maldonado, 2020). Though some crab spiders can increase the number of potential pollinators approaching the flower using deceptive signalling that exploit an insect’s ability to perceive UV colouration (Heiling et al., 2003; Llandres and Rodríguez-Gironés, 2011; Vieira et al., 2017), several studies have shown that the presence of a spider on a flower deters pollinators (Robertson and Maguire, 2005; Yokoi and Fujisaki, 2008). Clearly, some pollinators are capable of detecting these spiders (Defrize et al., 2010; Reader et al., 2006) and minimise their risk by evaluating the flower before landing (Ings and Chittka, 2008). What is not known, especially in natural conditions, is if insects can respond to a predation risk by altering their flight trajectories before landing and whether motion of the flowers affects their evaluation.

In this study, we evaluated the predator detection strategies of dune wasps (*Microbembex nigrifrons*; Hymenoptera: Crabronidae) as they approached a spider occupied *Palafoxia lindenii* flowerhead under two conditions of wind induced movement, i.e., when the flower was still and when it was moving. We measured the appearance of the flower and spider using psychophysical visual modelling from the perspective of a hymenopteran visual system and related the appearance to changes in wasp flight trajectories and landing decisions. If a wasp can detect the presence of the spider, we expected that their flight characteristics would reflect it and that the wasp would be able to detect spiders more easily when the flowers were still.

## METHODS

### Study site and species

The study was conducted in February and May 2019 at the Centro de Investigaciones Costeras La Mancha (CICOLMA), situated on the coast of the Gulf of Mexico (19° 36’ N and 96° 22’ W). *Palafoxia lindenii* A. Gray (Asteraceae) is an endemic dune pioneer species with white and purple inflorescences (hereafter ‘flowerhead’; (Álvarez-Molina et al., 2013). *Mecaphesa dubia* (Araneae: Thomisidae) is a colour-polymorphic spider that is frequently found on these flowerheads, either on top or to the side (i.e., on the receptacle). The most frequent colour morphs were white and purple (Rodríguez-Morales et al., 2018). *Microbembex nigrifrons* (Hymenoptera: Crabronidae) nests in dunes near vegetation (Evans et al., 2009), and feed on the pollen and nectar of *P. lindenii* flowers. These wasps are known to use visual cues to locate their nests and adult wasps provision their nests with dead arthropods (Alcock and Ryan, 1973).

### Visual Modelling

We quantified the visual appearance of the spiders as perceived by the wasp visual system at different distances using multispectral standardized images of female *Mecaphesa dubia* (n = 8) spiders positioned on the side and the top of the *Palafoxia lindenii* flowerhead. To do so, we took photos of the spider in both parts of the flowerhead using an Olympus Pen E-PM2 camera (converted to full-spectrum) with a UV transmitting EL-Nikkor 80 mm f/5.6 lens attached. We took two type of photos: one using a Baader Venus UV pass and the other with UV/IR cut filters, to obtain images in the ultraviolet (~ 300–400 nm) and in the human visible part of the spectrum (~ 400–700 nm) respectively. Each photo included two reflectance standards of 93% and 7% as well and a scale, and for a light source, we used an Iwasaki EYE Color arc lamp (70 W 1.0 A; Venture Lighting Europe Ltd., Hertfordshire, UK) with the UV block filter manually removed. The photos were saved in RAW format and processed using the Multispectral Image Calibration and Analysis (MICA) toolbox (Troscianko and Stevens, 2015) for ImageJ (v 1.52a), resulting in multispectral files with reflectance values of the spiders in the different position of the flowerhead.

Each multispectral file was converted to quantum catch files used in an integrative analysis by means of Quantitative Pattern Colour Analysis (QCPA) of the MICA toolbox. This is a framework based on the Receptor Noise Limited (RNL) model (Vorobyev and Osorio, 1998) and includes spectral sensitivity and visual acuity (Cronin et al., 2014). Since there is no information about the visual system of *M. nigrifrons* with respect to the visual traits mentioned above, we created a model wasp visual system using the data available for closely related species.

Thus, for the colour vision we used a reconstruction of the spectral sensitivity of *Philanthus triangulum* Fabricius (Sphecidae) with sensitivity peaks at: UV = 344, SW = 444, and MW = 524 (Peitsch et al., 1992). For the RNL model, we used the Weber fraction (w= 0.13) and the relative density for each receptor class (1:0.471:4.412 ratios for the UV:SW:MW receptors, respectively) based on the honeybee vision (Defrize et al., 2010). To model the visual acuity, we used the minimal resolvable angle value reported for *Bembix palmata* Smith (Crabronidae) equal to 1.22 cycles per degree (Feller et al., 2021).

We used the *‘Colour Maps’* approach to represent and delimit the entire visual scene combining visual acuity and spectral sensitivity data in a perceptually calibrated Hymenopteran trichromatic colour space (van den Berg et al., 2019). We estimated the portion of overlap between the spider and the flowerhead as perceived by the bee. The higher the overlap, the less likely it is for the viewer to perceive differences between the spider and the flowerhead. Finally, to have a representation of the noise reduction subsequent to visual acuity correction (Ligon et al., 2018) and recovering chromatic and luminance edge information, we included a RNL Ranked Filter based false colour image which is a representation of the colours using the wasp visual system we created. However, due to a lack of behavioural validation of the detection thresholds in the wasp, we use this image for visualisation purposes only. We compared the perceptual overlap of the colour maps generated for the spider and the flowerhead at different distances, in each position, by means of a Kruskal-Wallis test with a *Post-Hoc* Wilcoxon test done in the R statistics package.

### Flight trajectories of *M. nigrifrons*

We filmed wasps approaching *P. lindenii* flowerheads with a high-speed camera (Chronos 1.4, 500 fps, Krontech.com) with two wind treatments (Moving or Still) and the following conditions: 1. Flowerheads without spiders (Control, Moving: n = 9; Still, n = 6), 2. Flowerheads with live spiders tethered on the side (Side, Moving: n = 6; Still, n = 8, Fig. 1A), 3. Flowerheads with live spiders tethered on the top (Top, Moving: n = 8; Still, n = 6, Fig. 1B). Spiders were tethered to the flowerheads by means of a dental floss strand affixed to their ventral side with a non-toxic glue and placed into position with a pin. We stabilized flower stalks with a stick for the Still treatment and used freely moving stalks for the Moving treatment. This ensured that the constant sea breeze generated flower movement in the Moving treatment, but we did not control for extent of movement. The mean fluctuation of the flowers, as measured by the distance between the centre of the flower between frames, was significantly lower (Mann Whitney test statistic = 78, p < 0.0001) in the Still treatment in comparison with the Moving treatment. The cameras were placed at a distance of 1 m above the flowerheads, with the focal flower always centred; and the number of flowerheads per plant were similar. From the videos, the position of the centre of the flowerhead, head and abdomen of the wasp were manually tracked in 2D, using the MTrackJ plugin in ImageJ (Meijering et al., 2012). We recorded the following variables: Decision distance (i.e., the point where the wasp decides to land on or avoid the flower; this was recorded by visual inspection by one observer of the high-speed video in a frame-by-frame manner; see the red dot in Fig. 1 for an example), Inspection time (duration of flower observation, i.e., zig-zag flight, at close range), Outcome (whether the wasp avoided the flower or landed). From the coordinates of wasp position, we calculated the following metrics: Sinuosity index (a measure of deviation from a straight line) using the R package *trajr* (McLean and Skowron Volponi, 2018), the wasp’s body axis angle with respect to the flower (0° implies that the wasp was pointing directly at the flower) and speed.

**Fig 1:**
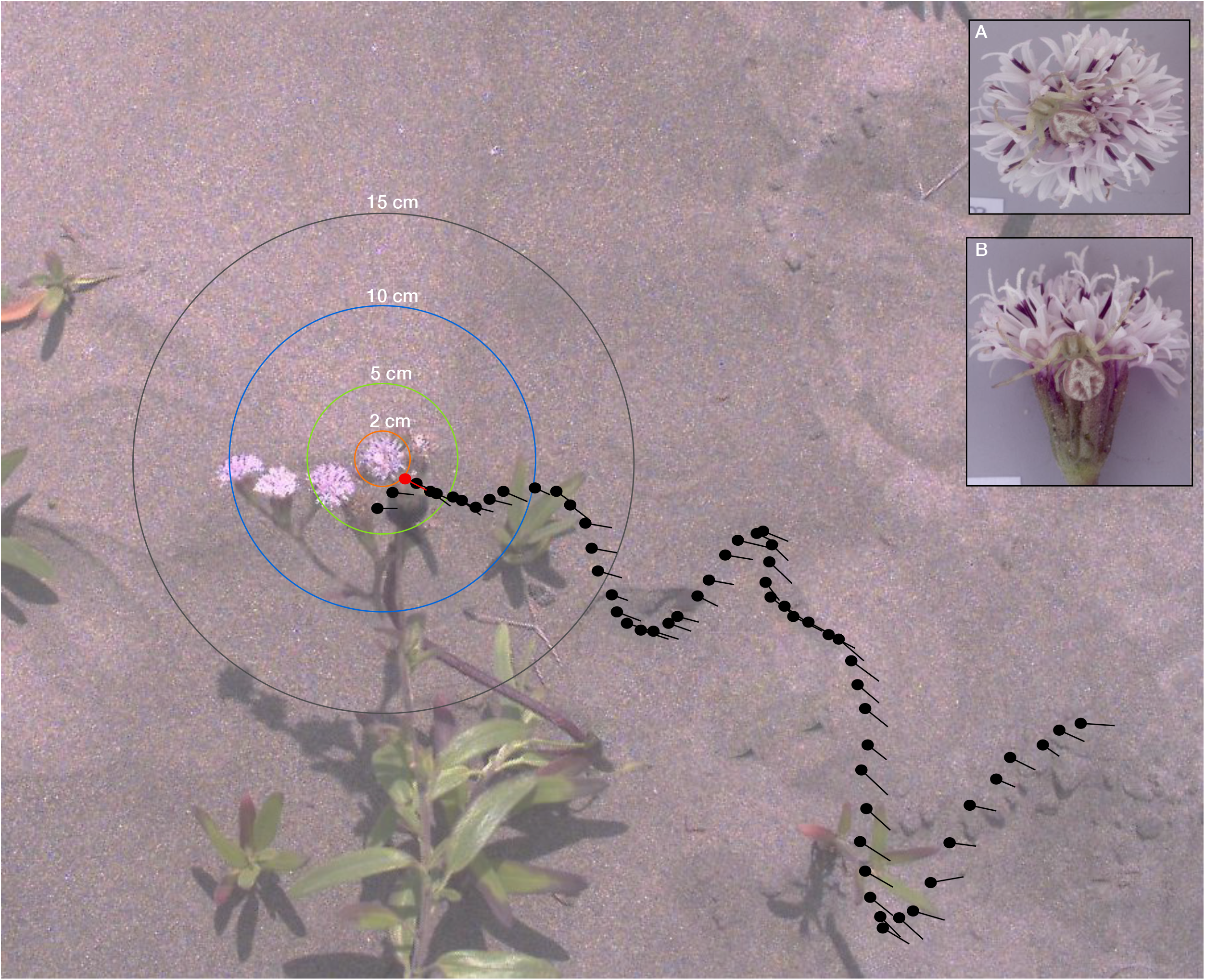
Sample trajectory of a *Microbembex nigrifrons* wasp approaching an unoccupied *Palafoxia lindenii* flowerhead as seen from the top. Wasp positions are subsampled for clarity. The point when the wasp made a decision (in this case to avoid the flower) is highlighted in red. Circles around the flowerhead represent the different distances used in the visual modelling analysis. The insets **A** and **B** show the *Mecaphesa dubia* spiders tethered above and to the side of the flowerhead respectively.

Distance profile curves, i.e., distance between the wasp’s position and the flower’s position at every point of the trajectory, were compared with an unsupervised cluster analysis based on the Dynamic Time Warping distance method which retains shape information (Fu et al., 2008; Hu et al., 2013; Keogh and Ratanamahatana, 2005) using Mathematica ver. 12. In order to compensate for differences in length of trajectories (i.e., some wasps flew for a longer time than others), we interpolated them such that all trajectories were of the same length. Trajectory data were coloured according to the distance to the flower, identifying four circular regions (at 2 cm, 5 cm, 10 cm, >10 cm) centred at the flower’s position (Fig. 1). We then analysed the frequency of distance profile shapes with respect to the different treatments with contingency tests.

To determine the direction of flight, we generated a line extending from the wasp’s abdomen and head (in the direction of the head; i.e., body axis). We subsequently compared the body axis angles of wasps approaching still and moving spider-occupied flowers with a Watson-Wheeler Two sample test using the *circular* package in R (Pewsey et al., 2013).

## RESULTS

### Visual modelling

We generated pseudo-colour images of the spiders on the flowerhead (Fig. 2A) that took into account both spectral sensitivity as well as visual acuity. The log Receptor Noise Limited (RNL) cluster modelling of the chromatic distances (ΔS), perceptual thresholds (1 Just-Noticeable Difference), and visual acuity of the wasp visual system showed that spiders may be detected only at a distance of around 5 cm from the flowerhead (Fig. 2B).

**Fig 2:**
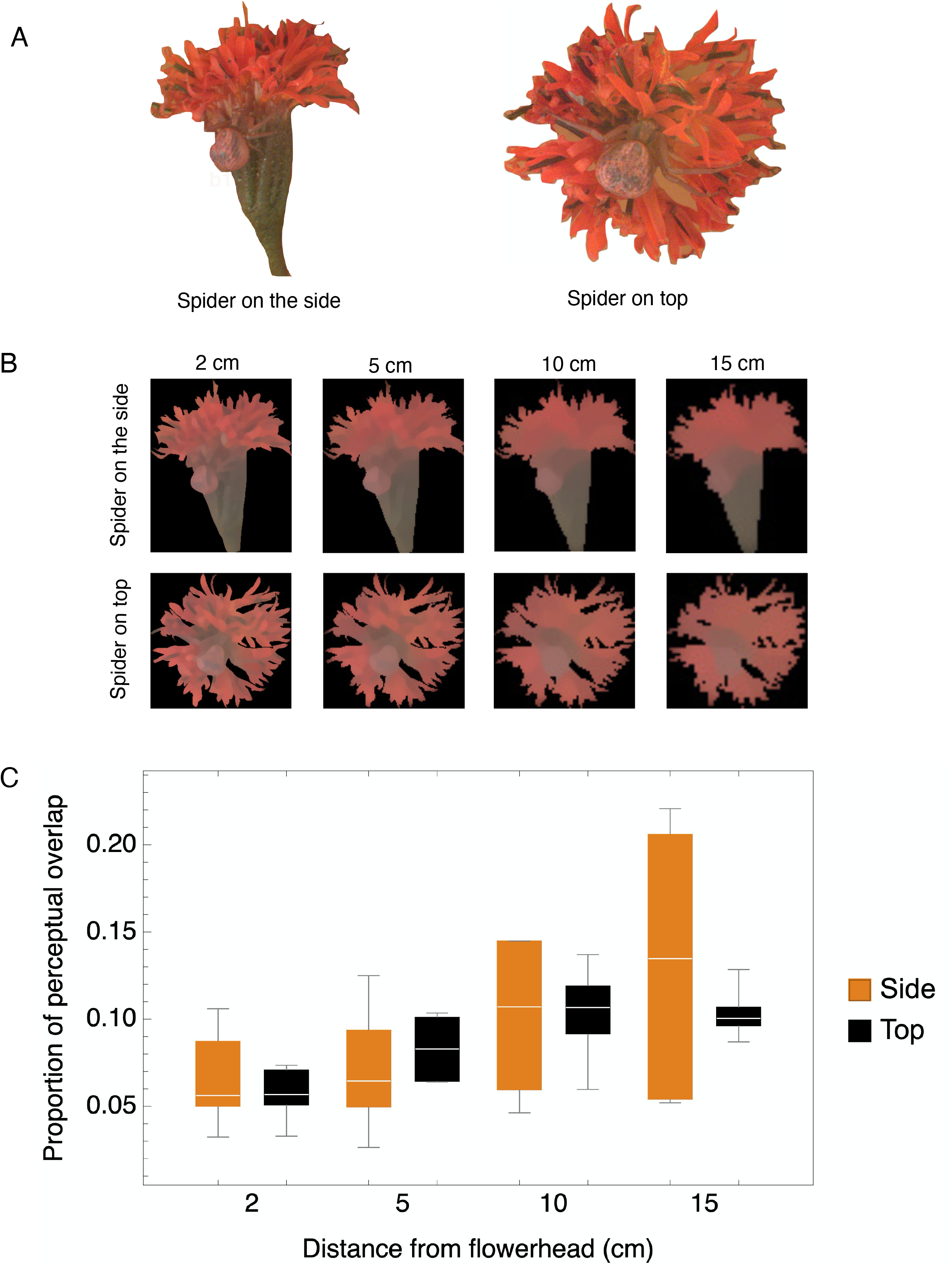
Colour modelling analysis summary of the Hymenopteran visual perception of *Mecaphesa dubia* spiders when located in the side or in the top part of the *Palafoxia lindenii* flowerhead at different observation distances (2, 5, 10, 15 cm). (A) False colour image simulating the perception of the wasp visual system. The image of the spider in the different parts of the flower were created for visualization purposes by assigning the colour blue, green and red for the UV, SW, and MW photoreceptor, respectively. (B) These panels show the results of a Receptor Noise Limited filter method, which performs noise reduction after the acuity control based on Gaussian filters while preserving chromatic and luminance edges, simulating spectral sensitivity and visual acuity of the wasp visual system at different distances of the spider located in the different parts of the flowerhead. (C) Using the wasp visual system created, we estimated the proportion of perceptual overlap between the spider and the flowerhead in the Hymenopteran colour space (higher overlap implies that it is more difficult for the viewer to perceive differences).

When comparing the overlap of the colour maps (Fig. S2) that represent the perception of the spider on the flowerhead by the wasp visual system with respect to the spider position, we found that the perceptual overlap is higher when the spider is on the top of the flowerhead (F = 19.7, df = 1; p < 0.001) and at a larger distance away from the flowerhead (F = 5.23, df = 3; p = 0.004; Fig. 2C). When the overlap is higher, the wasp would find it harder to visually separate the spider from the flower background. The interaction between position and distance was not significant. Thus, the perceptual overlap was significantly higher between 2 cm and 15 (p = 0.014) or 10 cm (p = 0.004), while there is no difference between the other pairwise comparisons.

### Wasp behaviour

Dune wasps approached flowers in a typical zig-zag flight (Fig. 3; see S1 for a video and S3 for a plot of all trajectories). Wasps were significantly more likely to land on flowers that were moving, irrespective of the presence of a spider (GLM, Logit Link, χ^2^ = 9.1, df = 1,37, p = 0.003; Fig. 4A) whereas spider location (χ^2^ = 3.2, df = 2,37, p = 0.193) and the interaction (χ^2^ = 1.3, df = 2,37, p = 0.512) were not significant. There was no significant effect of spider or wind treatment on the distance at which they made a decision to avoid or land on the flower (GLM, Identity link function, Gamma distribution, Spider: χ^2^ = 0.0006, df = 1,38, p = 0.97; Movement: χ^2^ = 1.39, df = 1,38, p = 0.24; Interaction: χ^2^ = 0.17, df = 1,38, p = 0.67; Moving: Mean ± S.D. = 5.14 ± 2.91 cm, Still: 6.45 ± 3.16 cm, Fig. 4B). However, there seems to be a trend that shows that wasps make their decisions earlier when approaching moving flowers. There was no effect of spider location (ANOVA: χ^2^ = 2.77, df = 2,37, p = 0.25), movement (χ^2^ = 0.54, df = 1,37, p = 0.463) or the interaction (χ^2^ = 1.99, df = 2,37, p = 0.36) on inspection time.

**Fig. 3:**
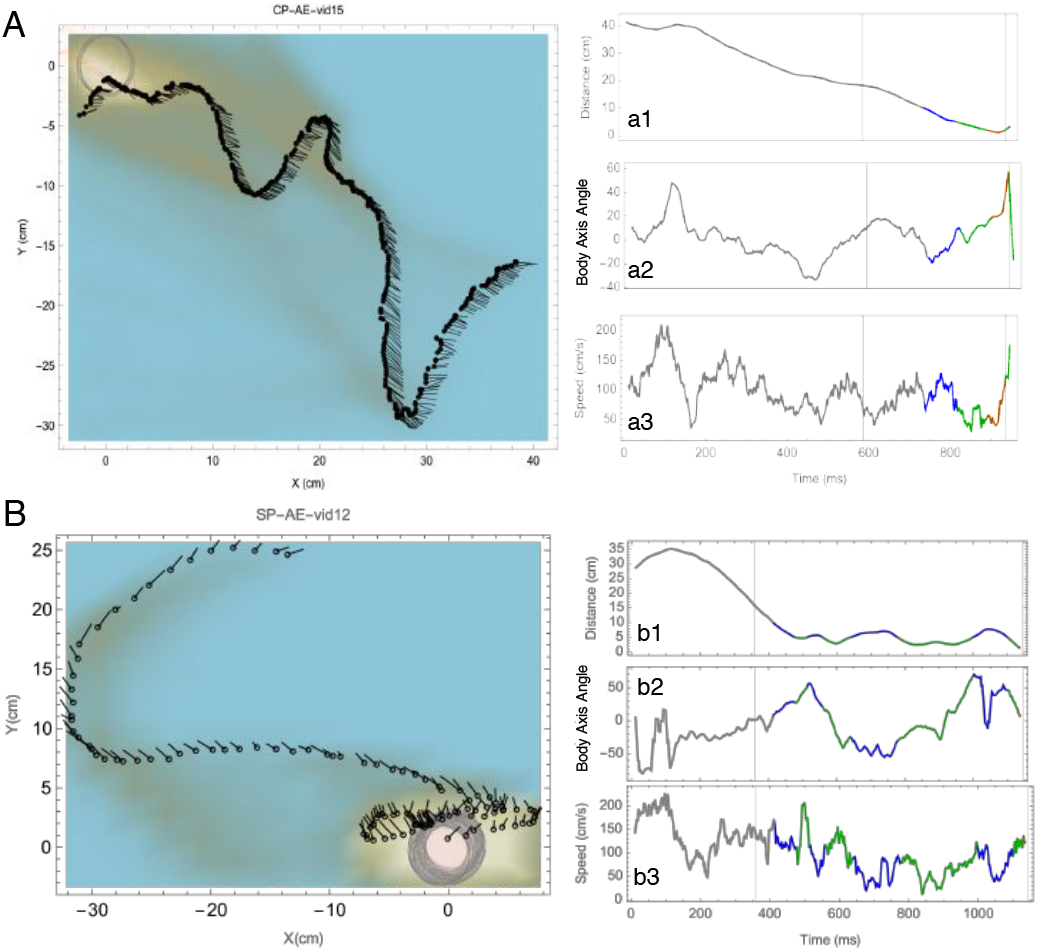
Sample trajectories in two movement treatments: A: Still and B: Moving, showing body axes in orange lines. Grey circle shows the extent of flower movement. Time series of Distance profile (a1, b1), Speed (a2, b2) and Body axis angle (a3, b3) of the sample trajectories. All curves are colour coded according to the proximity to the flower with Grey (> 10 cm), Blue (10-5 cm), Green (5-2 cm) and Orange (< 2 cm)

**Fig 4A:**
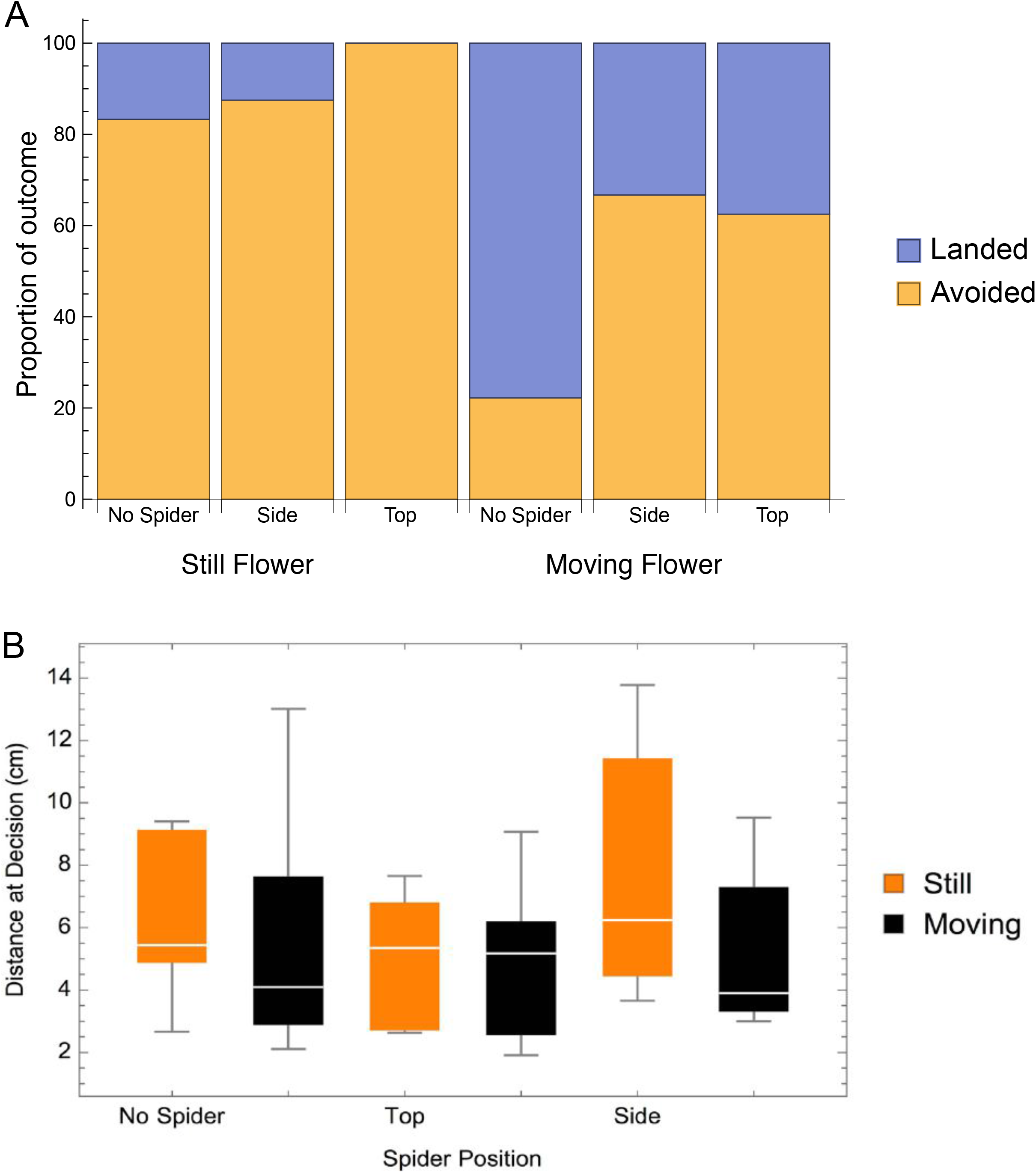
Frequencies of wasps that landed on or avoided flowerheads. Moving flowers with spiders were more likely to be visited that still flowers with spiders. 4B: The distances at which wasps decided to avoid or land at the flower.

### Distance profiles

Spider location, flower movement and their interaction significantly influenced wasp distance profile sinuosity (GLM, Identity link function, Spider: χ^2^ = 7.75, df = 2,37, p = 0.021; Movement: χ^2^ = 10.87, df = 1,37, p < 0.001; Interaction: χ^2^ = 7, df = 2,37, p = 0.030). Sinuosity was higher when the spider was on the side of the flowerhead in the still treatment, suggesting that wasps could respond to the presence of the spider in this condition.

The unsupervised DTW cluster analysis showed that the distance profiles (see Fig. 3a1, 3b1 for examples) of all wasp trajectories separated into two main clusters (designated accordingly as *Sinuous* and *Straight*; Fig. 5, see Fig. S4 for the distance matrix of the dendrogram). Contingency analysis of the frequency of type of distance profile showed that wasps had a straighter profile while approaching flowers without spiders and moving flowers with spiders (Fisher’s Exact Test; p = 0.012; Fig. 5 inset).

**Fig. 5:**
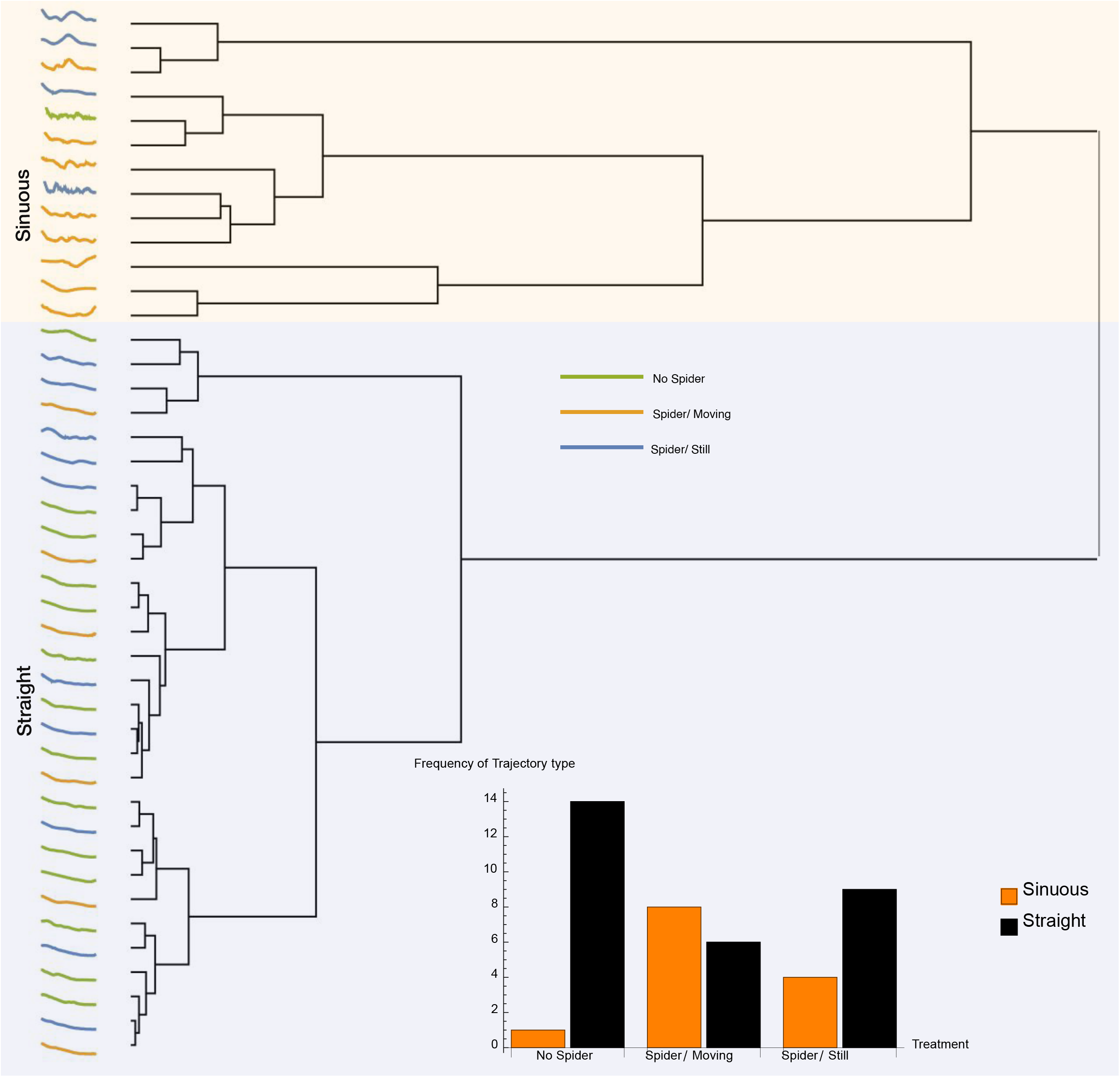
Results of a Dynamic Time Warping cluster analysis of distance profiles (a proxy for the trajectory shape) of all wasps approaching flowerheads with no spider (green lines), a moving flower with a spider (orange lines) and a still flower with spider (blue lines). The frequencies of sinuous and straight profiles are shown in the inset. Wasps approaching moving flowers with spiders were more likely to show a straighter distance profile.

### Body axis

Wasps consistently maintained a body axis angle centred on the flower (e.g., Fig. 3a2, 3b2, see orange lines in Fig. 3A,B) in both treatments (Still angle in radians: Mean ± S.D = 0.0066 ± 0.50, Rayleigh Test of uniformity = 0.88, p < 0.001; Moving angle in radians: Mean ± S.D = −0.034 ± 0.53, Rayleigh Test of uniformity = 0.86, p < 0.001). Wasps approaching moving spider-occupied flowers were more likely to maintain their body axis angle on the flower location with higher peaks in the angle histogram (Watson’s Two-Sample Test of Homogeneity, Test statistic = 0.5872, p < 0.001; Fig. 6).

**Fig. 6:**
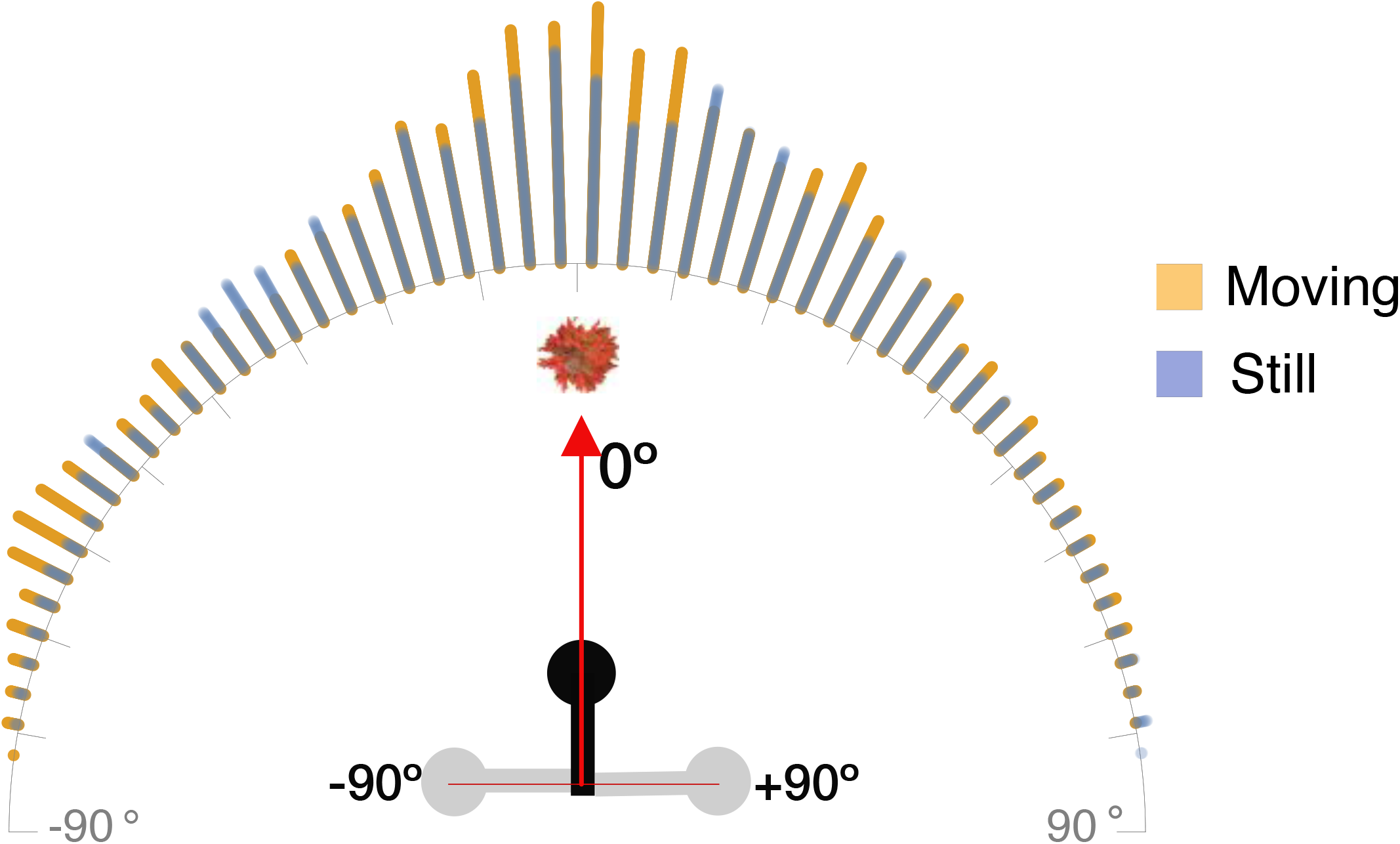
Frequency distribution of all wasp body axis angles as they approached moving (orange bars) or still (blue bars) flower heads with spiders. 0 degrees implies that the wasp’s body axis angle coincided with the flowerhead. Note that since wasps never turned around, the maximum extent of the body axis angle was always lesser than ± 90°. Body axis angle distributions was significantly different between the two conditions, with higher peaks when wasps approached the moving flowerheads with spiders.

### Speed

Wasps slowed down as they approached the flower (e.g., Fig. 3 a3, b3), and the average slowdown in speed was significantly different between the wind treatments (Fig. 7A). Wasps approaching moving flowers reduced their speed at a steeper slope than those approaching still flowers (Linear Regression: R^2^ = 0.828, F_1,1999_ = 9613.74, p = 0.046). Contrast post hoc analysis showed that wasps approaching still flowers were significantly faster than those approaching moving flowers at >10, 10 and 5 cm distances, whereas wasps approaching still flowers were significantly slower only at the 2 cm distance (ANOVA, df = 7, F_2,1999_ = 4809, p < 0.0001; Fig. 7B).

**Fig. 7A:**
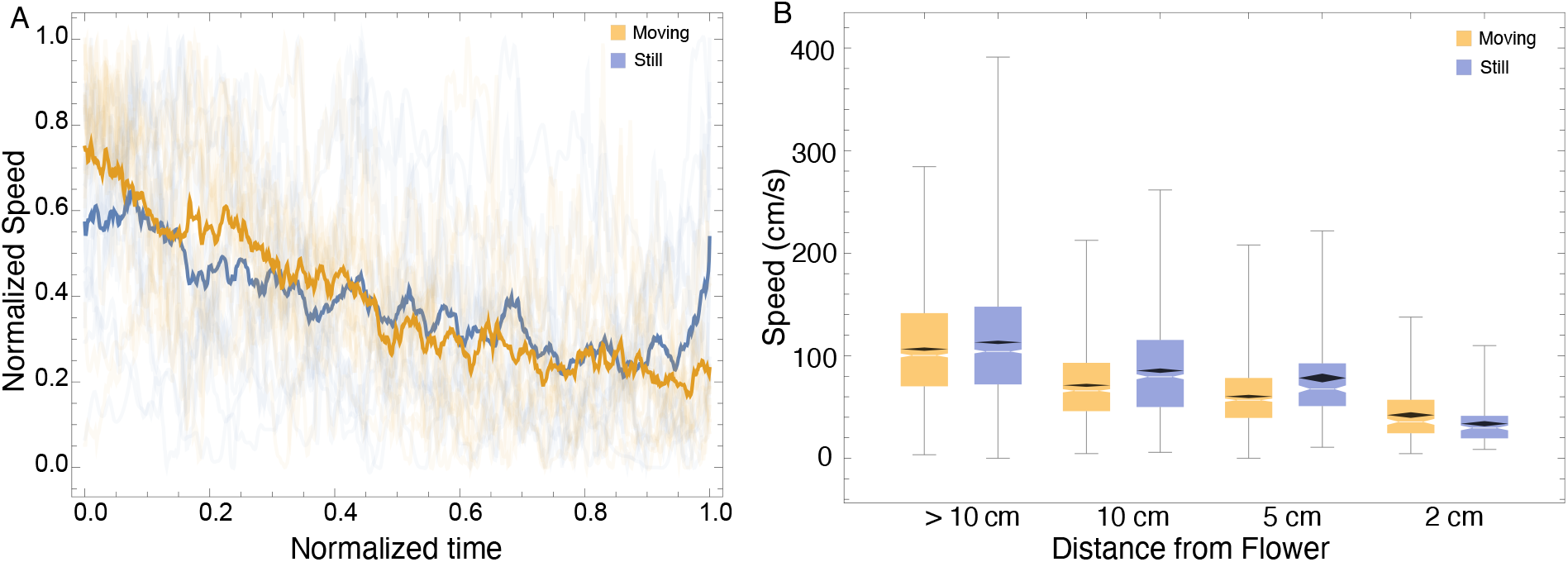
Normalised speed profiles of all wasps that approached moving (orange lines) and still (blue lines) flowerheads with spiders. The mean speed is shown with thicker lines. Wasps slowed down at significantly different rates as they approached the flowerheads. 7B: Boxplots showing median speed (notches) and mean confidence interval diamonds at different distance categories. Speeds were significantly greater in wasps approaching still flowers with spiders at the >10, 10 and 5 cm categories, whereas speed was greater in wasps approaching moving flowers at the 2 cm category.

## DISCUSSION

Dune wasps evaluate flowers for crab spider predators based on colour and motion cues, and their flight characteristics reflect their decision-making process. We found that though dune wasps locate and approach the flower from a distance, their decisions to land or avoid the flower occurs at a very close distance and this is due to the constraints of their perceptual system. When approaching moving flowers, wasps were more likely to land, had a straighter distance profile, steeper speed reduction and higher peaks in their body axis angle distribution targeting the flower.

Flight trajectories of insects are a useful window into their decision making process and can be used to understand their response to predation risk (Ings et al., 2011; Robertson and Maguire, 2005). For example, bumblebees that were trained to approach artificial flowers with cryptic or conspicuous artificial spiders showed a decrease in flight speed and an increase in inspection time after their first encounter with an attack (Ings et al., 2011). Bees were equally likely to avoid cryptic or conspicuous flowers, but their inspection times were longer for cryptic spiders (Ings et al., 2011). Wasp flight has so far been studied extensively, but largely from the perspective of learning flights, i.e., how a wasp learns the position of its nest (Collett and Zeil, 1996; Stürzl et al., 2016; Zeil et al., 1996) and comparatively little information is available about foraging decisions in free flying wasps. Wasps probably use the motion of the typical zig-zag flight to acquire visual depth information of the object in question (Egelhaaf, 2012; Lehrer, 1996; Lehrer and Campan, 2004). In hoverflies, the zig zag flight (termed ‘hesitation behaviour’) was more often seen when they approached flowers with a dead spider, and the authors attribute this to wasp flight mimicry, but it is more likely that this flight pattern allows the insect to gather more visual information regarding the predator (Nityananda et al., 2014; Yokoi and Fujisaki, 2008) and the motion parallax generated which would also aid the insect in range estimation (Kral and Poteser, 1997).

We expected that wasps would detect the presence of the spider at a larger distance when approaching still flowers. We did not find a statistically significant effect but in general the trend suggests that wasp do have difficulty in evaluating moving flowers. However, our study was based on 2D trajectories and a 3D reconstruction of flight trajectories should give better measurements of distances. Furthermore, the fact that wasps responded to the presence of the spiders at a very close distance suggests there might be multimodal processes at play here, such as chemical detection. In an experiment looking at honeybee response to spider occupied flowers, it was shown that bees were less likely to land on flowers that had been previously been exposed to spider cues, even when the spider was not present at the time of approach (Reader et al., 2006). The spider’s crypsis may be overcome by chemical detection, but since there is likely to be substantial variation in odour plume range and strength in natural conditions, a multimodal detection strategy is essential.

From the point of view of the spiders, there is an extensive literature on the effect of crab spider crypsis on potential pollinators (Dukas, 2019). The main lines of thought are as follows (*sensu* Brechbuhl et al. (2010)): 1. Spiders are cryptic to prey, 2. They are cryptic to predators, 3. They attract prey due to deceptive ssignalling, or 4. They can be detected and avoided by prey (i.e., no effect of crypsis). There is evidence for and against all four hypotheses, but using different crab spider species and different prey types. This variation in insect response to crab spiders (Rodríguez-Morales et al., 2019) can be attributed to the diversity in perceptual systems of the insects themselves. One would expect that if the main prey is of hymenopteran origin, then there would be a higher likelihood of successful evasion of predation in comparison to other insects that do not perform an inspection behaviour such as certain flies. Our study emphasises the need to take a species centric approach (*sensu* von Uexhill’s *umwelt* (Caves et al., 2019; Uexküll, 2013)) to understand these interactions at a fine scale.

One of the unexpected results from our study was the relatively low frequency of wasp landings on unoccupied still flowers. We suggest that wasps are evaluating flowers, using both visual and olfactory cues (which we did not test), perhaps for traces of earlier visits by conspecifics or predators. Evaluating flowers for predation risk is significantly influenced by flower motion, suggesting that the cognitive processes needed to integrate all this information is compromised under certain abiotic conditions (Nityananda et al., 2014).

Our study shows that prey response to predators occurs at fine scales and the prey’s perceptual biases play a significant role in assessing risk. To avoid an attack, wasps need to detect the predator at a close range and then respond by manoeuvring out of range before the attack occurs in order to maximise their escape rate.

## ACKNOWLEDGEMENTS

We thank Derian Jesus Jaime Duran and Rogelio Rosales García for logistical help, Ajay Narendra and Diana Perez Staples for helpful comments on the manuscript and Armando Falcón Brindis for the taxonomic determination of the wasp. We thank the Instituto de Ecología (INECOL) for use of the CICOLMA facilities. This project was financed by a Ciencia Basica grant (CB-2016-01-285529) from CONACyT, Mexico to DR.

## AUTHOR CONTRIBUTIONS

Data Collection - DRM, LRO

Writing Original Draft - DR

Writing & Revision- DR, DRM, HTM, LRO

Analysis - DR, DRM, HTM, LRO

Visualisation - DR, HTM, LRO

Funding Acquisition - DR

Conceptualisation - DR, DRM

Project Administration DR, DRM

## Supplementary files

S1: A sample video showing a wasp approaching an unoccupied still flowerhead.

S2: Colour maps that show the degree of overlap between the spider (a1), the flowerhead (f1) using the Hymenopteran colour space. See text for details.

S3: All trajectories of wasp approaches to Moving (orange lines) and Still (blue lines) flowers. Note that the flowerhead is for illustrative purposes and not to scale.

S4: Distance matrix of the Dynamic Time Warp based unsupervised classification of distance profile of wasp trajectories.

